# Remotely-sensed tornado signatures indicate an ecological threshold for eagle nest destruction

**DOI:** 10.1101/049387

**Authors:** Jeremy D. Ross, Cheryl L. Cavert, Lena C. Larsson

**Affiliations:** George Miksch Sutton Avian Research Center, Oklahoma Biological Survey, University of Oklahoma, P.O. Box 2007, Bartlesville, OK 74005 USA; Bald Eagle Survey Team (BEST), P.O. Box 2007, Bartlesville, OK 74005 USA

**Keywords:** severe storm ecology, disturbance, nest failure, environmental stochasticity, catastrophe

## Abstract

Bald Eagles (*Haliaeetus leucocephalus*) will reuse massive nests placed high in trees over multiple years, potentially exposing them to catastrophe loss during severe storms. The stochasticity of localized weather, however, has traditionally been viewed as impeding the quantitative study of such risks. In March 2015 a severe weather outbreak along the Arkansas River near Tulsa, Oklahoma caused widespread damage among a highly-concentrated and long-monitored population of nesting Bald Eagles. We conducted field surveys on the extent of nest loss and consulted weather and observer records to determine what characteristics of the storm (maximum azimuthal wind shear) or nests (years of use) might have been associated with nest destruction. We found 5 of 9 nests along a ~24km stretch of the river were destroyed during the storm, causing the death of at least 8 eaglets. Mean years of use was higher among destroyed nests (4.8) than surviving nests (4.0), though not significantly so within this limited sample. The degree of maximum azimuthal shear (i.e., wind rotation) during the storm within 800m of the nests, however, did significantly differ both in terms of maximums (15.8 vs 9.5 ms^−1^) and means (9.4 vs 6.6 ms^−1^) for destroyed versus persisting nests, respectively. Our findings suggest a threshold of tornadic wind shear beyond which Bald Eagle nests, irrespective of age, could be prone to catastrophe. Such insights are key to developing accurate models of population persistence, especially in light of potential shifts in severe weather patterns under various climate change scenarios.

## Introduction

Stochastic events such as severe weather are difficult to predict or model with respect to anticipated spatiotemporal impacts, especially far in advance and for localized phenomena like tornadoes. The devastating forces that comprise tornadoes represent acute disturbances on the landscape that are mostly limited to a few hundred meters wide and tens of kilometer-long paths (1), although there are obviously much-publicized catastrophic tornadoes (2). With this in mind, population viability modeling incorporating stochastic weather impacts has largely relegated these to random variables with “best-estimate” parameters (3,4). Yet, there exist many decades of reported accounts of storm damage (5), developing global networks of biologists and citizen scientists (6,7), and emerging technology with the capacity to accurately quantify storm characteristics and relate those to effects on the ground (8–10). There are likely ways to more accurately parameterize the stochasticity of tornadoes and other rare but devastating events, with important ecological and conservation ramifications (11).

Relatively fine-scale severe weather data has become increasingly available as hardware and software technology has advanced. Modern Doppler weather radars (e.g., WSR-88R or “NEXRAD” units used in the USA) provide a measure of radial velocity of objects relative to the transceiver that can be used to estimate the movement of objects aloft, such as wind-blown raindrops. Doppler shifts within adjacent radial range gates of the radar data (i.e., airspace voxels) can also serve to inform estimates of localized wind shear. In fact, such data form the basis for remotely-detecting mesocyclone formation in supercell thunderstorms in near real-time and is used by the National Weather Service to inform the issuing tornado warnings (12). Advances in available software used to analyze NEXRAD data have provided ground-truthed ways to estimate the severity of two thunderstorm forces in particular: a) the maximum estimated size of hail (MESH) (13,14) and b) the maximum azimuthal wind shear (15). The latter measure is most relevant to the quantification of cyclonic wind forces and could be central to refining models of biological impacts during tornado outbreaks.

During the late afternoon and evening of March 25, 2015 a tornado was spawned during a severe thunderstorm outbreak over Sand Springs in western Tulsa and southeastern Osage counties, Oklahoma. It cut a path of structural and biological damage along at least a 17.9km path and, at its largest, was rated as an EF-2 tornado with an estimated width of 732m and maximum estimated wind speeds of 201-217km/h (16). Impacts of the storm included human casualties (1 killed, 30 injured), numerous homes and buildings damaged or outright destroyed, and widespread uprooting and limb damage among mature hardwood trees (16). The stretch of the Arkansas River affected by the storm damage also hosted the second highest concentration of nesting Bald Eagles (*Haliaeetus leucocephalus*) in the state of Oklahoma, as determined by ongoing monitoring efforts of the George Miksch Sutton Avian Research Center (“Sutton Center”) and its citizen-scientist Bald Eagle Survey Team (BEST).

During the 20+ years since self-sustaining breeding populations of Bald Eagles were successfully re-established in Oklahoma, the Sutton Center has monitored all known nests of the species in the state in order to track productivity. Along a 24km length of the Arkansas River in western Tulsa County there were nine active nests within 1km of the main channel during the spring of 2015 (Figure 1). The number of successive years that each nest had been used was known, as was the status of each nest within 7 days before and after the March 25^th^ storm.

**Figure 1:**
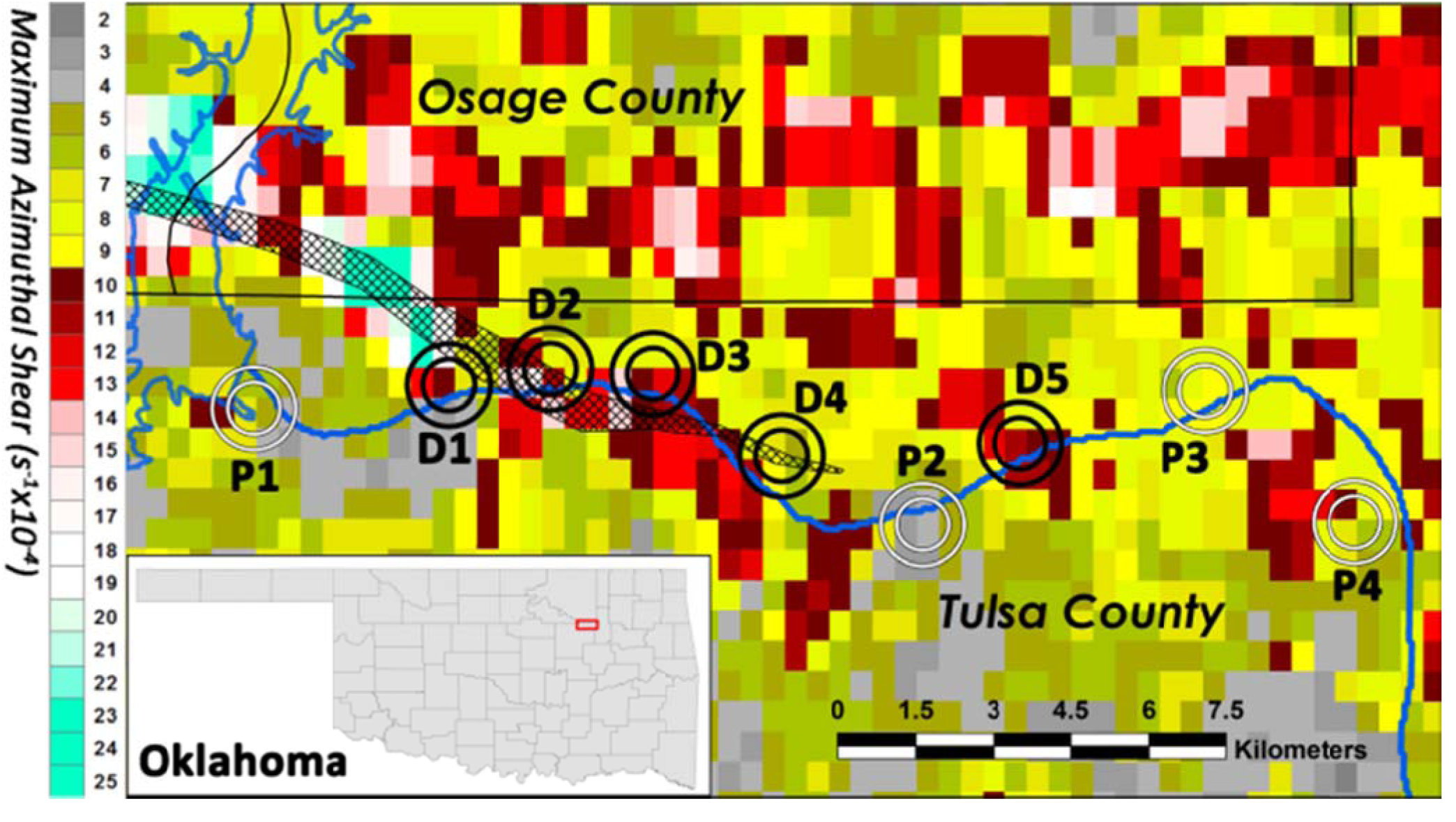
Map of WDSS-II maximum azimuthal shear associated with the March 25, 2015 Sand Springs tornado relative to 500m and 800m distances from the nine Bald Eagle nests which were alternately destroyed (black rings) or persisted (whiterings) during the storm. Shown is the area of focus within the inset (red box atop map of Oklahoma), the estimated tornado path (crosshatched area; NWS Tulsa 2015), county boundaries (black lines), and the Arkansas River including the Keystone Lake impoundment (dark blue line).

After observing that various Bald Eagle nests of approximately similar size, composition, and height in trees had alternately been destroyed or persisted through the tornadic storm, we recognized the tragic opportunity to examine the correlation between severe storm characteristics and biological impacts on the ground. To our knowledge, this would be the first attempt to quantitatively assess the association between measurable characteristics of tornadic winds and their immediate impacts on animals.

Others have documented tornado impacts on birds, yet most either studied space-use shifts within the avian community long after the tornado had passed (17,18) or provided anecdotal reports of detectable damage without comparative control data or sufficient sampling (19–24). Johnson (25) did conduct two transects of point counts adjacent to and through farmland areas struck by a tornadic hailstorm, finding an apparent reduction in birdlife as well as shifts in the avian community as a result of the damage. However, as Johnson admitted, his surveys came eight days after the storm and his delineation of the impacted area was subjectively-defined. Wiedenfeld & Wiedenfeld (26) likewise conducted quantitative post-storm surveys for evidence of bird mortality, in their case two days after a tornadic front caused widespread mortality among Neotropical migrants that were likely approaching the Louisiana Gulf Coast during their migratory crossing. Though the authors were able to extrapolate their results to very large impacts (i.e., tens of thousands of birds killed), they lacked the capacity to directly link the tornado to the losses. Such widespread avian mortality could have alternatively been related to a frontal shifting of winds, a situation known to lead to widespread drowning among migratory birds(27–29).

The ubiquity and diversity of biology as a possible source of post-tornado data provides the rare but widespread opportunities needed to assess severe weather forces. Examining single-species effects, as we do in this study, can allow subsequent extrapolation across the species range and better predictions of possible future impacts. Better parameterization of stochastic events in ecological models, especially within an increasingly changing and variable climate, could be central to predicting the long-term viability of populations.

## Materials and Methods

### Field Surveys

As part of the regular nest monitoring program conducted by the BEST citizen-scientists, each active Bald Eagle nest in the Sand Springs, OK area was monitored at an undisruptive distance (>250m) approximately every other week during the breeding season (mid-January through early June; all assigned to C. Cavert). At each visit we noted evidence of apparent incubation, adult visitations, and estimated age of chicks (if visible) in order to track the active status and eventual productivity of each nest. In mid-March 2015, prior to the nest trees leafing out, we took a reference photo for each of the nests. Resident Bald Eagles in Oklahoma will remain territorial year-round, but yearly challenges from other adults can lead to relatively high turnover (30). Therefore, we assigned each active nest to the territory within which it was located. Even in the case of adult turnover, the history of each discrete nest in terms of number of consecutive years occupied was known.

In the aftermath of the tornado of March 25^th^, 2015, we revisited all potentially-impacted Bald Eagle nest sites being monitored in the impacted area as soon as emergency road closures were lifted (mostly by the next day, with all but one site visited within five days). During these checks we assessed whether the nest had been destroyed or persisted, if there was evidence of eaglet deaths, the direction destroyed nests had been blown (if possible among the debris), and we took another reference photo from the same vantage point used for photos taken prior to the storm.

### Weather Radar Data

We downloaded data pertaining to the weather characteristics concurrent with the Sand Springs tornado formation from the National Severe Storm Laboratory’s warning decision support system-integrated information (WDSS-II)(31) using the portal available at http://ondemand.nssl.noaa.gov/. The WDSS-II system incorporates relative velocity data across all available elevational sweeps of NEXRAD data and runs these through a linear least squares derivative filter (15) to calculate an azimuthal shear field (in per-second units). The maximum value of shear across multiple radars and across the lower 3km of the atmosphere are then calculated to a grid of pixels measuring 5.0x10^−3^ degrees for each latitude and longitude, and are stored as raster files which can be exported to a GIS mapping program.

From the WDSS-II portal we downloaded the georeferenced azimuthal shear data for the entire area potentially impacted by the Sand Springs storm, selecting a broad time span (17:00-21:00 on March 25, 2015) which fully encompassed the period of tornado formation through dissipation (i.e., 17:21-17:38)(16). These data were imported into ArcGIS (ESRI, Redlands, CA) and, because they are provided in a simple RGB raster format, assigned pixel values for maximum azimuthal shear according to the WDSS-II color scale. In order to provide enough pixel data to provide a suitable quantification of shear within proximities of the nests, we resampled the data using a nearest-neighbor algorithm into blocks measuring 5.0x10^−4^ degrees for each latitude and longitude. This produced an overlay identical to the original data, though with 100X more pixels to allow sufficient quantitative comparisons among the airspaces surrounding the Bald Eagle nests.

Since we were interested in determining if the radar-estimated shear values were significantly predictive of localized destruction on the ground, we independently evaluated shear within two radial distances from each Bald Eagle nest: 500m & 800m. We specifically determined the maximum, mean, median, and standard deviation for azimuthal shear value within these airspace columns for each of the nine nests along our focal stretch of the Arkansas River. We then used a t-test to independently evaluate whether the number of years of use significantly differed between nests which persisted through the storm and those which were destroyed. Finally, we conducted an ANOVA for each possible distance and summary data set (e.g., maximum within 800m) with age of nest as a cofactor.

## Results

Field surveys of nests within the affected stretch of the Arkansas River west of Tulsa, OK revealed that 5 of 9 Bald Eagle nests active during the 2015 breeding season had been destroyed during the Sand Springs tornado event (Figures 1 &2). All destroyed nests were located north of the river, whereas three of four surviving nests were located south of the river. The mean age of nests among those destroyed during the storm (4.8 years) was higher than the nests which persisted (4.0 years), although the difference was not significant between these limited samples (p=0.599; Table 1).

**Table 1.** Azimuthal wind shear characteristics of the March 25, 2015 Sand Springs tornado near nine Bald Eagle nests that alternately persisted through the storm or were destroyed during the event.

| | | | Azimuthal wind shear metrics in proximity to nest ($\text{ms}^{-1}$) | | | | | | | | |
| --- | --- | --- | --- | --- | --- | --- | --- | --- | --- | --- | --- |
| Map |  | Fate | Years Active | Within 500m |  |  |  | Within 800m |  |  |  |
| Location | Nest Nickname |  |  | Max | Mean | Median | Std Dev | Max | Mean | Median | Std Dev |
| D1 | Tanglewood | destroyed | 3 | 23.0 | 8.6 | 10.0 | 3.7 | 23.0 | 8.8 | 7.0 | 5.2 |
| D2 | Wekiwa | destroyed | 7 | 16.0 | 10.7 | 11.0 | 2.9 | 16.0 | 10.5 | 11.0 | 3.2 |
| D3 | 152<sup>nd</sup> W Ave | destroyed | 5 | 15.0 | 10.1 | 10.0 | 1.9 | 15.0 | 10.1 | 10.0 | 2.4 |
| D4 | Rader | destroyed | 6 | 11.0 | 7.1 | 7.0 | 1.6 | 13.0 | 7.6 | 7.0 | 1.7 |
| D5 | Hot Rod | destroyed | 3 | 12.0 | 10.5 | 11.0 | 1.2 | 12.0 | 10.1 | 11.0 | 1.3 |
| P1 | White Water Pk | persisted | 1 | 10.0 | 6.3 | 6.0 | 1.4 | 10.0 | 6.3 | 6.0 | 1.8 |
| P2 | Avery Drive | persisted | 5 | 5.0 | 3.7 | 4.0 | 0.6 | 7.0 | 4.1 | 4.0 | 1.2 |
| P3 | Berryhill Creek | persisted | 7 | 9.0 | 8.6 | 9.0 | 0.6 | 9.0 | 8.2 | 8.0 | 0.9 |
| P4 | H-F Refinery | persisted | 3 | 12.0 | 7.8 | 7.0 | 1.6 | 12.0 | 8.0 | 8.0 | 1.9 |
| *Mean (destroyed)* |  |  | 4.8 | 15.4 | 9.4 | 9.8 | 2.3 | 15.8 | 9.4 | 9.2 | 2.8 |
| *Mean (persisted)* |  |  | 4.0 | 9.0 | 6.6 | 6.5 | 1.1 | 9.5 | 6.6 | 6.5 | 1.4 |
| *p-value\** |  |  | 0.599 | 0.059 | 0.078 | **0.048** | 0.092 | **0.041** | **0.047** | 0.108 | 0.157 |

\*based on t-value for Years Active, F-value for all others

**Figure 2:**
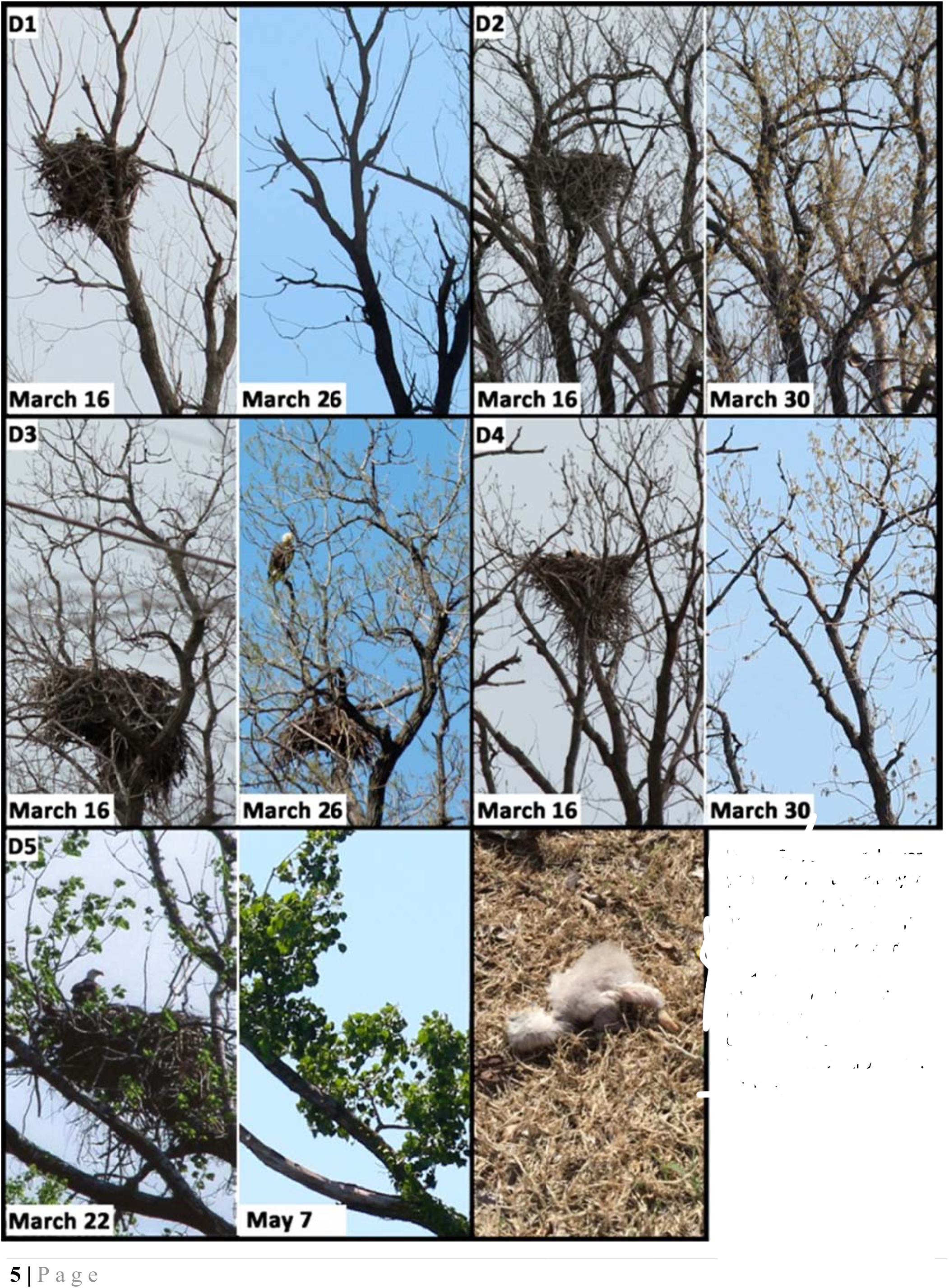
Before and after photos of nests destroyed during the March 25, 2015 tornado (C. Cavert). Nest labels correspond to locations indicated in Figure 1. The bottom right panel shows a 10-d old eaglet that died when nest D4 was destroyed (C. Gomez).

Except at the 152^nd^ W Ave site (D3 in Figure 1) nest destruction constituted complete obliteration, often with little other damage to the nesting tree limbs (Figure 2). At only two of the five destroyed nests could we unequivocally determine the direction the nest was blown from its tree: Tanglewood (D1) was blown southward, while Rader (D4) was blown northeastward. Through direct observation, reports from wildlife agents, or consultations with local residents, we were able to determine that 8 to 10 young eaglets ranging in age from approximately 10 days to three weeks old died as a result of their nests being destroyed during the tornado (Figure 2). Two of the four nests which persisted through the storm, two went on to successfully fledge at least one eaglet (P1 & P2), one failed by the middle of April (P4), and one had actually failed and been abandoned days prior to the tornado but still represented an intact nest (P3; Figure 1).

Azimuthal wind shear varied widely across the area of study, especially among the Bald Eagle nest sites of focus (Figure 1). After accounting for differences in nest age, the maximum, mean, and standard deviation of shear forces (in units of ms-^1^) within 500m of the nests were higher among destroyed nests (max=15.4, 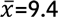; s.d.=2.3) versus those which persisted (max=9.0, 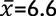; s.d.=1.1), though not quite at levels of significance (p-values: 0.059, 0.078, & 0.092, respectively; Table 1). The median shear values around nests, however, were significantly higher among destroyed nests (9.8) than among nests which persisted (6.5; p=0.032; Table 1). Within 800m of the nest sites the maximum and mean shear forces were both significantly higher among destroyed nests (max=15.8, 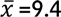) versus nests which persisted (max=9.5; 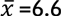; p-values: 0.033 and 0.032, respectively), though median and standard deviations did not significantly differ (p-values: 0.083 and 0.140, respectively; Table 1).Age of nests was not a significant factor in any of the ANOVA examined (range: F-values= 0.046-0.845; p=0.393-0.837).

## Discussion

Among the most iconic visuals associated with the United States of America are the Bald Eagle and large, devastating tornadoes. During the noteworthy instances when these phenomena overlap, there are likely ecological ramifications that could have lasting impacts on the long-term conservation of Bald Eagle populations. The analyses presented in this study offer the chance to better parameterize what may be population-limiting events for Bald Eagles and other species (11,32), especially in light of increasing climate instability which could lead to more frequent, widespread, and powerful convective storms (33–35).

It appears that a threshold might exist for Bald Eagle nest persistence versus destruction during periods of cyclonic wind shear. In our case, the most striking difference associated with nest destruction was the maximum wind shear value within 800m; the possible tipping point at azimuthal shear of 12ms^−1^. All examples of destroyed nests experienced a maximal radar-estimated shear within 800m at or above this threshold, while maximum shear within 800m of persisting nests fell at or below this level (Table 1). This level of wind shear exceeds the absolute maximum forces estimated for ~25% of 80 tornadoes evaluated by Visco et al. (36) which had formed over the contiguous United States. In other words, ~75% of all tornadoes in the USA may produce azimuthal shear in excess of the theoretical level required to destroy a Bald Eagle nest.

The H-F Refinery nest (P4) appears to have withstood the highest proximate wind shear forces during the March 25^th^, 2015 storm (Table 1). There are two interrelated reasons why it may have persisted: 1) the angle of approach of the tornadic storm, and 2) the nest location relative to the riverbank. Cyclonic winds comprising tornadoes in Oklahoma rotate in a counter-clockwise fashion. The brunt of the winds would have approached the P4 nest across a highly disrupted industrial development (i.e., an oil refinery), and this collection of tall and rigid structures could have served to disrupt the winds experienced at ground level. This is a stark contrast to the riverside nests which would have experienced largely-unimpeded winds swirling across the open water (Figure 1). Our observations suggested that nest D1 was thrown southward by the approaching tornado, while nest D4 was blown to the northeast and away from the river when the tornado skirted along its northern bank (16).

## Measuring global severe weather threats for animals

Tornadoes represent the most powerful aggregations of force within the atmosphere in terms of energy per unit area (1). It is, therefore, unsurprising that their occurrence leads to devastation among the biological community. The unpredictability, relatively acute impacts, and debris fields associated with tornadoes can make them a challenge to study. However, like any complex natural phenomenon, severe weather impacts could be quantified with vigilance, communication, and readiness among potential observers. Such data may prove useful in cobbling together a better understanding of severe weather effects, plus how to best model and predict such outcomes.

Gathering data on biological impacts attributable to tornadoes is challenging but, as the present study demonstrates, is possible if the damage coincides with ongoing field monitoring programs. The value of using damage to bird life as a metric of tornadic strength was perhaps surprisingly given scientific consideration more than 170 years ago. In a bizarre fashion, Elias Loomis in 1842 fired a freshly-killed chicken from a cannon using a small charge in an attempt to simulate the potential windspeeds of a strong tornado. His intention was to test anecdotal assertions that domestic fowl are plucked clean during such violent winds (37). Though this seems absurd (it won the 1997 Ig Nobel Prize for meteorology), reports of tornado-plucked chickens are sufficiently frequent that Loomis’ approach was not completely misguided (38).

Among the published examples of tornado damage to birdlife, some estimates place the minimum estimates for the numbers killed during a single event in the thousands (19) or even tens of thousands (22,25,26). The effects of tornadic damage also apply to other vertebrate and invertebrate animal species. For example, developmental disruption in the form of asymmetrical growth was reported in the aftermath of a EF-3 tornado for both mice (39) and spiders (40) occupying a damaged portion of an Ohio forest. Stress during a severe hailstorm associated with a tornadic supercell also appeared to affect the development of ground-nesting sparrows in Oklahoma, leaving lasting signatures within growing feathers (41).

Provided that post-storm surveys can sufficiently account for scavenging (e.g., by arriving soon after the storm, or at least measuring evidence of past and ongoing carcass removal) as well as detectability of carcasses, the potentially large sample of storm casualties across the animal community can easily provide the opportunity to geographically plot the estimated impacts. Such scientifically-defensible survey data would be necessary to properly assess storm impacts relative to other georeferenced datasets, such as remotely-sensed landscape features or climatological data.

The advent of modern meteorological equipment has enabled scientists to quantify relatively high-resolution spatiotemporal data of severe storm forces. This technology has provided the means to not only better understand how weather forms, but has also been co-opted for biological purposes since it can detect animals aloft just as easily as water droplets (42). In addition to the azimuthal wind shear and MESH products available through WDSS-II, there are other emerging weather analysis products that could prove useful for severe storm ecology studies—particularly from dual-polarization radars (43). Even data from satellite-based sensors can provide great insight into severe storm ecological processes at large scales. For example, Klimowski et al. (44) were able to quantify from *GOES-8* data the vegetative damage from a tornadic hailstorm across a 120km path in western South Dakota. Rudimentary field surveys confirmed the widespread plant damage, although no data was collected which could specifically compare the apparent gradient of destruction or estimate the immediate and secondary impacts on the local animal community.

Beyond the acute effects of tornadoes, the high winds associated with gust fronts, derechos, and downbursts can lead to widespread and substantial impacts for tree-nesting raptors. For instance, Gilmer & Stewart (45,46) found that over a three-year period 37% and 34% of all Ferruginous Hawk (*Buteo regalis*) and Swainson’s Hawk (*B.swainsoni*) nest losses, respectively, could be attributed to severe storms producing high winds and hail over their North Dakota study site. Roth & Marzluff (47) reported a similar rate of nest failure attributable to hail-producing windstorms among *B. regalis* nesting in Kansas (31.3%). Wintering raptors can likewise face severe windstorm risks; Bernitz (48) found hundreds of migratory kestrels (*Falco spp*.) dead or severely injured after a favored roost tree in South Africa partly collapsed during high winds. In fact, many bird species are potentially exposed to the damaging effect of storm-generated winds: heron rookeries may be partly or completely destroyed (21,24,49,50), floating nests of grebes can be swamped or obliterated (51), and resident cavity nesters may be permanently displaced after fleeing the storm (52).

In addition to the five Bald Eagle nests lost in Sand Springs, OK on March 25, 2015, a nest further to the north near Lake Yahola was also destroyed despite being many kilometers from the tornado’s path. A possible culprit was a sudden downburst near the hail core of the storm (16). Such forces can be extremely powerful (1) and, since that nest fell directly downward and nearly intact, the downburst could have been sufficient to snap the limb supporting the Bald Eagle nest.Unfortunately, downbursts are not as easily quantified from NEXRAD data (15), so an examination of this potential destructive force was not conducted.

## Severe weather ecology under climate change

The destructive forces of severe storms arise from the thermal energy available to fuel convective uplift, as well as the vertical wind shear most often associated with the jet stream (1). The resulting kinetics of updraft and rotation within convective storms produces tornadoes, severe hail, heavy downpours, lightning, and occasional downbursts when the uplift collapses. Because climate change scenarios are in disagreement regarding the predicted changes in the strength of updrafts and vertical wind shear, the anticipated changes in severe storm intensity, frequency, and spread are not yet well defined (53). However, some analyses are pointing toward a likely acceleration of devastating extreme storm prevalence over the next century (33,35).

Shifts in severe storm timing and geographic spread could have profound implications for animal communities, particularly breeding birds. In areas which historically have not been routinely impacted by severe weather, the biological community may have poorly-evolved behavioral or physiological capacities to persist through extreme ecological disturbances (e.g., plants lack resistance adaptations in the areas where such events are rare) (54). Furthermore, where there may have historically been little overlap between severe weather timing and critical breeding periods, climate shifts could especially push migratory species increasingly toward a riskier spring arrival and nest initiation phenology, as has been shown among American White Pelicans breeding colonies in North Dakota (55). Provided that the magnitude of mass mortality events involving birds has significantly increased over the past 75 years (56), the emerging conflicts between biological life-cycles and climatic shifts may be having a global effect.

Southern populations of the Bald Eagle were once among the most recognizable examples of a species that had been reduced to a fraction of sustainable levels. In Oklahoma and surrounding states, breeding populations of this species were non-existent in the early 1980’s. Through a concerted effort, organizations such as the Sutton Center were able to recover these extirpated populations from translocated stock. If a catastrophic loss associated with a direct tornado strike had impacted these released populations at a critical time — such as when juveniles were being acclimated at hacking towers or when the first batch of releases were establishing their first nests — then the recovery of this threatened species could have been significantly set back. As evidenced by the way some nests appeared plucked from their trees with little damage otherwise (Figure 2), even moderately-strong storms pose a threat if they coincide with vulnerable stages of the life cycle. A similar threat certainly still faces other rare and declining species which occupy, even as migratory stopovers, those areas around the globe which are particularly susceptible to severe weather outbreaks.

## Acknowledgments

This study would not have been possible without the volunteer monitoring efforts of the citizen-scientists of the Sutton Avian Research Center’s Bald Eagle Survey Team (BEST). We also thank Tulsa County Game Warden Carlos Gomez for photos and information of nest damage in areas closed to the public. This study was funded by the Oklahoma Biological Survey.

